# Composite Biofidelity: Addressing Metric Degeneracy in Biomechanical Model Validation and Machine Learning Loss Design

**DOI:** 10.64898/2026.04.05.716563

**Authors:** Amruta Koshe, Ehsan Sobhani-Tehrani, Kian Jalaleddini, Hamid Motallebzadeh

**Author notes:** **Contributions:** HM designed and conceived the study; HM and AK performed the simulations; HM, AK, and EST performed the inference and analyses; EST and KJ contributed to the conceptualization of the idea; HM wrote the manuscript; all authors contributed to the manuscript.

## Abstract

Spectral similarity is often judged with a single metric such as RMSE, yet this can be misleading: physically different errors can produce similar scores. This is a critical limitation for computational biomechanics, where spectral agreement underpins both model validation and machine-learning loss design. Here, we develop a multi-metric framework for objective spectral biofidelity and test whether it better captures meaningful disagreement across complex frequency-domain responses.

We evaluated 12 complementary similarity metrics, including CORA and ISO/TS 18571, using controlled spectral perturbations that mimic common real-world deviations such as resonance shifts, localized spikes, and broadband tilts. We then applied the framework to an SBI-tuned finite-element middle-ear model to assess convergence with training dataset size and robustness to measurement noise across repeated stochastic runs.

No single metric performed reliably across all distortion types. Shape-based metrics tracked resonance morphology but could miss vertical scaling, whereas MaxError remained important for narrowband anomalies that smoother metrics underweighted. CORA and ISO 18571 did not consistently outperform simpler metrics. Rank aggregation using Borda count provided a robust consensus across metrics, enabling objective identification of training-data saturation and noise thresholds beyond which similarity rankings became unstable.

These results show that spectral biofidelity cannot be reduced to a single norm. A multi-metric consensus provides a clearer and more physically meaningful basis for comparing experimental and simulated spectra, and offers a more defensible foundation for data-fidelity terms in physics-informed and simulation-based machine learning.

## I. INTRODUCTION

Computational models are widely used to interpret complex biomechanical responses and to extrapolate beyond what can be measured experimentally (Motallebzadeh et al., 2018). Trust in these predictions depends on rigorous validation against experimental observations (Anderson et al., 2007; Oberkampf et al., 2002). Increasingly, this validation relies on ML-driven parameter tuning, including SBI methods that infer full posterior distributions (Cranmer et al., 2020; Motallebzadeh et al., 2025; Sackmann et al., 2019). This shift creates two linked needs: an objective definition of spectral agreement between computational and experimental data, and a loss function that reflects that agreement so optimization favors biofidelity over mathematically convenient minima.

The main challenge in objective validation is that spectral mismatch is multidimensional (Mottershead & Friswell, 1993; Terven et al., 2025). Experimental and simulated spectra can differ in overall level, resonance height and bandwidth, slope, phase, or the frequency location of key features, and these discrepancies do not carry the same physical meaning. Some spectral variation also arises from benign measurement factors, such as probe placement, boundary conditions, or reference pressure location, rather than true biomechanical change (Motallebzadeh et al., 2017, Figure 6). An objective comparison must therefore detect physically meaningful deviations while remaining tolerant to expected measurement-driven variation, especially when several independent spectra are evaluated together and a single similarity score can mask clinically or mechanistically important differences (Motallebzadeh et al., 2017; O’Connor et al., 2017).

In ML optimization, the loss function does more than score agreement; it determines what the model learns to reproduce and what it learns to ignore. If that loss does not reflect the aspects of agreement that are physically meaningful, optimization can converge to solutions that are less interpretable or even physically implausible, despite achieving lower numerical error (Goodfellow et al., 2016; Wang et al., 2004). This risk is especially pronounced for complex spectral responses, where distinct discrepancies in energy, shape, phase, or feature location can be traded against each other under a single scalar objective, producing “mathematically improved” fits that do not preserve the intended physical interpretation (Raue et al., 2009; Terven et al., 2025).

RMSE is widely used in model validation and ML training because it is simple, differentiable, and reduces point-to-point disagreement to a single scalar value. However, RMSE alone is often insufficient for rigorous comparison between experimental and simulated spectra and can be misleading for spectral biofidelity (Liemohn et al., 2021). Because RMSE emphasizes large-amplitude regions, it can favor fits that lower pointwise error while distorting important spectral features (Goodfellow et al., 2016; Mottershead & Friswell, 1993). This is especially problematic in frequency responses, where local resonances and phase trends often carry the main mechanistic information (Puria, 2003). Consequently, physically distinct distortions can produce similar RMSE values, creating metric degeneracy in which the algorithm achieves “better” scores while degrading the spectral features that define the underlying mechanics (Terven et al., 2025).

Composite rating tools such as CORA and ISO/TS 18571 were developed for time-domain waveform comparisons and are effective in that setting (International Organization for Standardization, 2014; Thunert, 2017). However, frequency-domain validation depends on mismatch patterns that are less directly captured by time-domain corridor scoring, including resonance distortions, broadband spectral tilt, and shifts in the frequencies of key spectral features. Composite objectives are also common in ML, for example in imaging and neural audio, where multiple loss terms are combined to capture different aspects of agreement (Johnson et al., 2016; Yang et al., 2021). In probabilistic model calibration, including SBI, this same issue remains: posterior inference still depends on how spectral agreement is scored, motivating a multi-metric framework for frequency-domain biofidelity (Motallebzadeh et al., 2025).

Existing objective biofidelity tools are largely time-domain or application-specific, and we are not aware of a widely adopted data-driven framework that integrates complementary similarity metrics for frequency-domain biofidelity across multiple spectra. This gap matters for both validation and ML-driven calibration, because metric-specific blind spots can produce degenerate agreement scores and unstable conclusions when several independent objectives must be matched at once. To address this, we use our SBI-tuned finite-element middle-ear model as a motivating case study and proceed in two phases. First, we systematically evaluate twelve similarity metrics across controlled spectral error topologies to identify the discrepancy patterns each metric detects or misses. Second, we introduce a Composite Biofidelity Framework that aggregates complementary metric classes across independent spectra to provide a single, physically defensible basis for multi-objective validation and parameter tuning. This perspective is also relevant to physics-informed ML, where the data-fidelity term should remain sensitive to physically distinct spectral distortions rather than optimize a single scalar loss. Multi-metric objectives therefore offer a principled route to more meaningful training targets across multiple outputs.

## II. METHODS

This section outlines our spectral similarity framework using an SBI-tuned finite-element middle-ear model as a case study. We compared experimental and simulated spectra for two purposes: to assess convergence as training dataset size increased, and to evaluate robustness as simulation noise was systematically varied.

### A. Reference Experimental Data

Experimental data were obtained from two independent measurement modalities in a single cadaveric temporal bone (specimen TB24). Wideband tympanometry was used to measure ear-canal input impedance and calculate acoustic absorbance, defined as the ratio of absorbed to input acoustic energy. Stapes velocity was also measured to characterize middle-ear acoustic transmission. To capture measurement and test-retest variability, seventeen independent sets of velocity, impedance, and absorbance spectra were collected at ambient pressure. These datasets served as both the physiological target for SBI and the reference data for metric evaluation (Motallebzadeh et al., 2025).

### B. Finite-Element Model Construction

A deterministic FE model of the middle ear was developed from post-experimental high-resolution μCT images of the same temporal bone (TB24). The reconstructed geometry was imported into COMSOL Multiphysics and included the tympanic membrane, ossicular chain, suspensory ligaments, and middle-ear cavity. Seven free parameters, including key joint stiffnesses, tissue moduli, and damping factors, were selected for inference based on their strong influence on system transfer functions and their clinical relevance to pathology detection. Prior values were sampled within physiologically plausible ranges to generate 10,000 frequency-domain training datasets simulating stapes velocity, impedance, and absorbance. Full model details are provided in (Motallebzadeh et al., 2025).

### C. Simulation-Based Inference (SBI) Framework

The SBI framework (Tejero-Cantero et al., 2020) was trained entirely on the pre-simulated FE datasets. It learned the relationship between randomly sampled prior parameter values and the corresponding acoustic and vibratory model outputs. When provided with the experimental reference data, the trained network returned posterior probability distributions for the seven free parameters. To generate the spectra used in the present similarity analyses, the FE model was run using the maximum a posteriori (MAP) parameter estimates and 10 samples drawn from the posteriors. Detailed network architectures and training hyperparameters are provided in (Motallebzadeh et al., 2025).

### D. Experimental Design: Learning-Curve and Noise-Effect Analyses

To assess how closely the SBI-inferred spectra matched the experimental reference data under different conditions, we performed two sensitivity analyses:

1. *Convergence (Learning-Curve) Analysis:* The SBI framework was trained on logarithmically spaced dataset sizes ranging from 100 to 10,000 simulations. For each training size, the composite biofidelity score was used to identify the convergence point beyond which additional data produced only marginal improvement.
2. *Noise-Robustness Analysis:* The training dataset size was fixed at 10,000 simulations. Gaussian noise, scaled from 0.1 to 10.0 times the baseline standard deviation of the experimental test-retest data, was added to the simulation outputs before SBI training to evaluate robustness to measurement and instrumentation noise.

Because SBI includes stochastic variation from network initialization, noise injection, and subset selection from the full training set, both analyses were repeated across 10 fixed random seeds. For each experimental condition, spectral similarity was evaluated using 11 simulated cases: the MAP estimate and 10 posterior samples. This design made the similarity estimates and rank aggregation robust to stochastic variation.

### E. Similarity Metrics

To capture the multidimensional nature of spectral disagreement, we evaluated 12 complementary similarity metrics. These metrics span four classes: magnitude and energy, shape and angular agreement, transport and morphological differences, and composite biofidelity standards. Each metric compares a reference spectrum *Y* with a simulated spectrum Ŷ sampled at *N* frequency points *f*. Some metrics are defined analytically, whereas others are computed algorithmically.

#### Magnitude and Energy Metrics

These metrics quantify pointwise amplitude differences and overall error magnitude. They are useful for bounding global disagreement, but are generally less sensitive to frequency shifts and spectral shape changes.

##### Root Mean Square Error (RMSE)

Standard L_2_ norm that penalizes larger deviations more strongly.

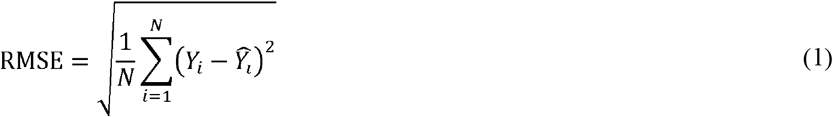

##### Mean Absolute Error (MAE)

Standard L_1_ norm that applies a linear penalty to absolute differences.

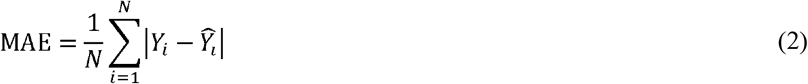

##### Maximum Error (MaxError)

Chebyshev distance, representing the single largest deviation across the spectrum.

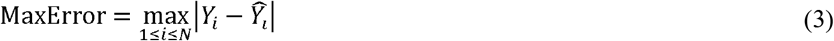

##### Weighted RMSE (wRMSE)

Frequency-weighted L norm designed to reflect the physiological and clinical importance of different frequency regions in middle-ear mechanics.

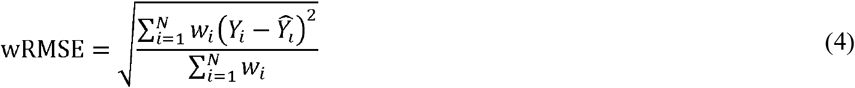

The weights w_i_ were assigned across five functional frequency bands, reflecting specific middle-ear mechanics (refer to Supplementary Information for details):

100–300 Hz (w = 0.15): stiffness-controlled region, sensitive to compliance-related changes such as negative middle-ear pressure or effusion.

300–800 Hz (w = 0.20): stiffness-to-inertance transition region, including fundamental frequencies and lower speech formants.

800–2,000 Hz (w = 0.25): primary resonance and peak power-transfer region, highly sensitive to conditions such as ossicular discontinuity.

2,000–5,000 Hz (w = 0.25): mass-controlled region, important for high-frequency consonant cues.

5,000–10,000 Hz (w = 0.15): extended-high-frequency region, strongly influenced by probe-depth artifacts and relevant to emerging clinical measures.

#### Shape and Angular Metrics

These metrics emphasize geometric similarity between spectra. They are sensitive to resonance shifts and spectral tilt, while being less sensitive to uniform vertical offsets.

##### Inverted Cosine Similarity (1 – Cosine)

Measures the angle between the two spectra treated as high-dimensional vectors, capturing shape similarity independent of overall amplitude.

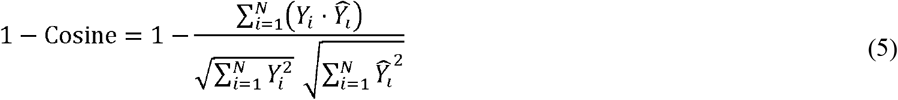

##### Inverted Pearson Correlation (1 – Pearson)

Measures linear shape similarity after mean-centering, making it sensitive to trends such as slope while ignoring absolute level.

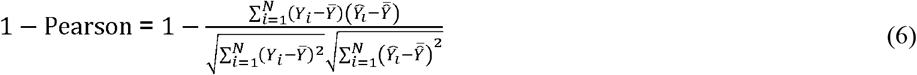

Where 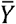 and 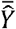 represent the mean amplitude of the entire experimental target spectrum and the entire simulated model spectrum, respectively.

#### Transport and Morphological Metrics

Unlike point-to-point error measures, these metrics evaluate how one spectrum must be transformed to match the other. They are therefore more tolerant of feature shifts and local warping along the frequency axis.

##### Wasserstein Distance (Earth Mover’s Distance)

Measures the minimum transport cost required to transform one distribution into the other, and is effective for tracking resonance shifts across frequency (implemented via scipy.stats).

##### Dynamic Time Warping (DTW)

Finds the optimal non-linear alignment between two spectra, allowing local compression or expansion along the frequency axis (implemented via dtw-python).

##### Energy Distance

Statistical distance between spectral energy profiles that satisfies the formal properties of a metric (implemented via scipy.stats).

##### Logarithmic Area Difference (AUC Diff)

Integrates absolute residual error over the logarithmic frequency axis so that broad high-frequency regions do not dominate localized low-frequency discrepancies.

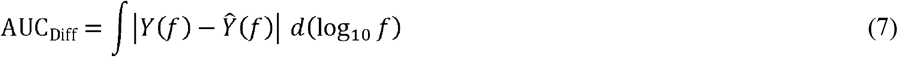

#### Composite Biofidelity Standards

To provide broader summary measures, we adapted two composite standards used in structural and impact biomechanics to quantify biofidelity, that is, how closely a surrogate reproduces the target biological response (Gehre et al., 2009). Custom Python implementations were developed from the published manuals.

##### CORA (Cross Correlation Analysis)

Combines a global Corridor method with a Cross-Correlation method. The Corridor score evaluates if the signal falls within a defined tolerance band, while the Cross-Correlation score evaluates the phase, magnitude (size), and shape characteristics. The final metric assigns a 50% weight to both primary components:

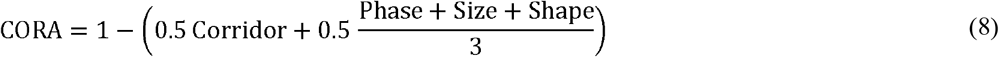

##### ISO/TS 18571

An objective rating metric that calculates a weighted global score by independently evaluating specific signal distortions. Following the standard specifications, the final metric was calculated using a 40% weighting for the corridor fit (utilizing 5% inner and 50% outer boundaries) and 20% weightings for phase, magnitude, and slope respectively:

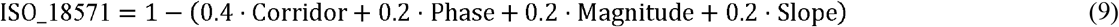

### F. Non-Parametric Rank Aggregation

We used a nonparametric rank-aggregation framework to compare simulated and experimental spectra across multiple similarity metrics and stochastic repeats, and to identify the condition with the strongest overall agreement. The procedure consisted of four steps:

1. *Spectrum generation:* For each of the 21 experimental conditions, and for each random seed, we generated 11 simulated spectra: one maximum a posteriori (MAP) estimate and 10 posterior samples.
2. *Metric evaluation:* For each condition, 12 similarity metrics were computed by comparing the 11 simulated spectra with the reference experimental spectra. Metric values were then averaged across the 11 spectra to yield one representative value per metric, per condition, per seed.
3. *Within-seed ranking:* Within each seed, the 21 conditions were ranked from best to worst for each metric. Ranks were then summed across the 12 metrics to obtain a raw Borda score for each condition.
4. *Across-seed aggregation:* Raw Borda scores were averaged across the 10 seeds and min-max normalized to a [0, 1] scale, where 0 indicates the most biofidelic condition.

### G. Convergence and Robustness Criteria

#### 1) Optimal Training Size (Saturation Point)

To identify the training size required for spectral convergence, we applied a geometric knee-point criterion to the aggregated Borda curve:

##### Monotonic envelope

To reduce stochastic fluctuations, Borda curves were first smoothed using a Savitzky-Golay filter and then converted to a monotonic non-increasing envelope using a cumulative minimum.

##### L-method knee point

The saturation point, marked with a red star in the figures, was defined as the point with the greatest perpendicular distance from this envelope to the secant line connecting the N=100 and N=10,000 endpoints.

#### 2) Measurement Noise Robustness

##### Normalization

Scores were min-max normalized from 0.0 at the noise-free baseline to 1.0 at the maximum degradation observed at the highest noise level.

##### 50% degradation threshold

The robustness limit was defined as the noise level at which the smoothed normalized curve crossed 0.5, corresponding to a 50% loss in biofidelity relative to the maximum observed degradation.

## III. RESULTS

### A. Evaluation of Metric Sensitivity and Degeneracy

#### Divergence of Individual Similarity Metrics

To evaluate the discriminative power and potential blind spots of the 12 similarity metrics, we applied twelve controlled spectral distortion scenarios (solid red) to a reference spectrum (dashed black) as shown in Fig. 1. The scenarios were designed to reflect common experimental and physiological perturbations, including calibration offsets (vertical shifts), probe or setup related feature shifts, smoothing and noise effects, and localized resonance irregularities that can arise from mass or stiffness changes.

**Fig. 1.**
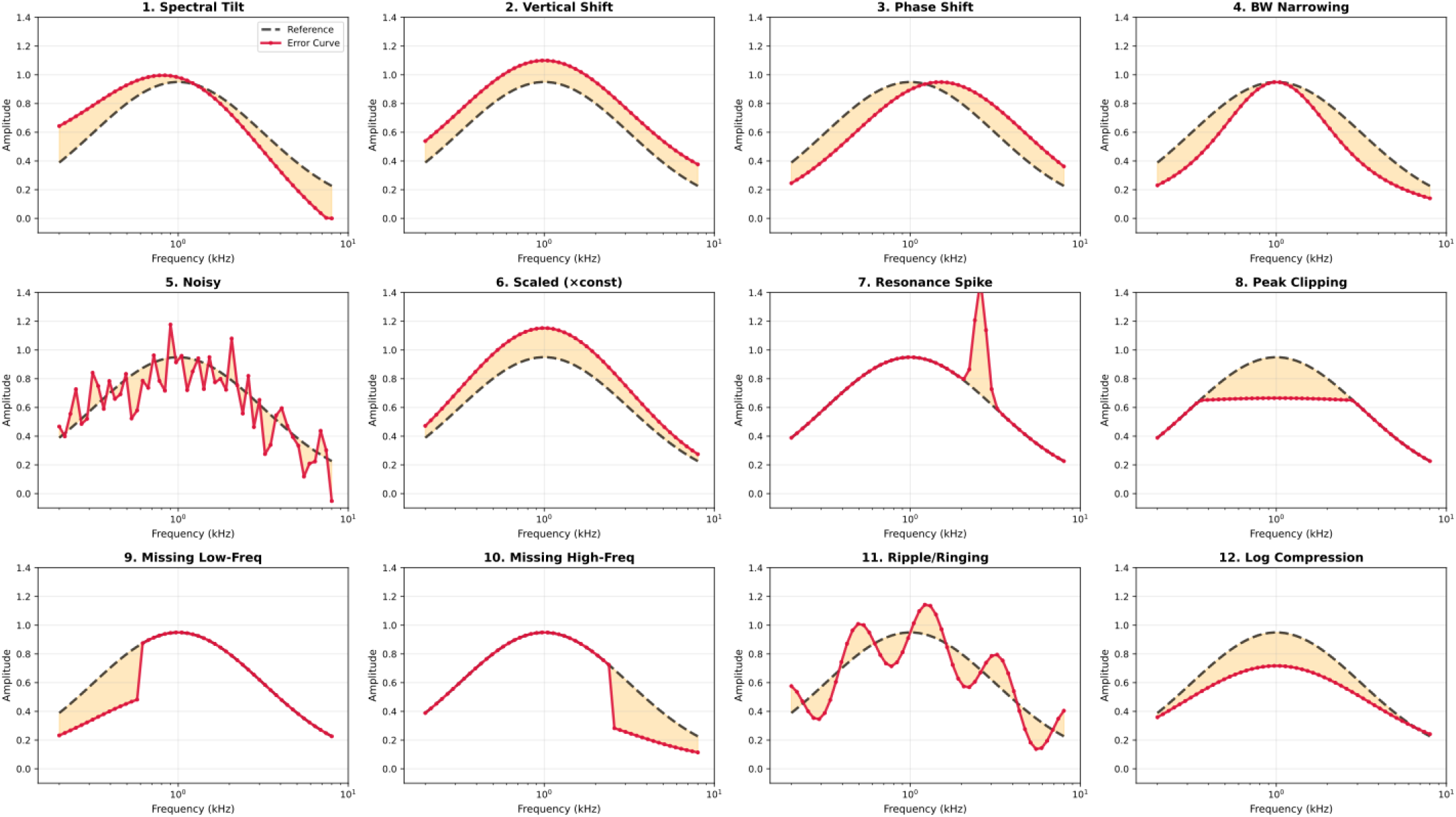
Simulated spectral error scenarios with constant RMSE. Twelve distinct spectral distortion scenarios (solid red) are applied to a reference spectrum (dashed black). The scenarios represent common perturbations, including calibration offsets, probe-or setup-related feature shifts, smoothing or noise effects, and localized resonance irregularities. All scenarios were scaled to the same RMSE (0.15), showing that visually distinct spectral distortions can share the same pointwise error score.

Each scenario was scaled using an optimization routine so that the total pointwise error energy was identical across cases (RMSE = 0.15). By holding RMSE constant, this comparison isolates sensitivity to spectral morphology and shows that distinct distortion patterns can receive the same RMSE, highlighting the need for multidimensional similarity assessment beyond a single pointwise energy metric.

#### Analysis of Metric Discrimination

To quantify how strongly each metric distinguished among the twelve iso-RMSE scenarios, we summarized its variability across scenarios using the coefficient of variation (CV), computed across the twelve cases. We then visualized scenario-specific metric responses in a heatmap of min-max normalized penalty scores, weighted by each metric’s CV so that metrics with greater across-scenario variability contributed proportionally more to the discrimination map (Fig. 2). In this representation, discrimination is reflected by the spread within each metric column rather than by the absolute color value itself. Nearly uniform columns indicate that a metric responds similarly across scenarios and therefore provides limited scenario-specific discrimination, whereas broader within-column spread indicates stronger selectivity across distortion types.

**Fig. 2.**
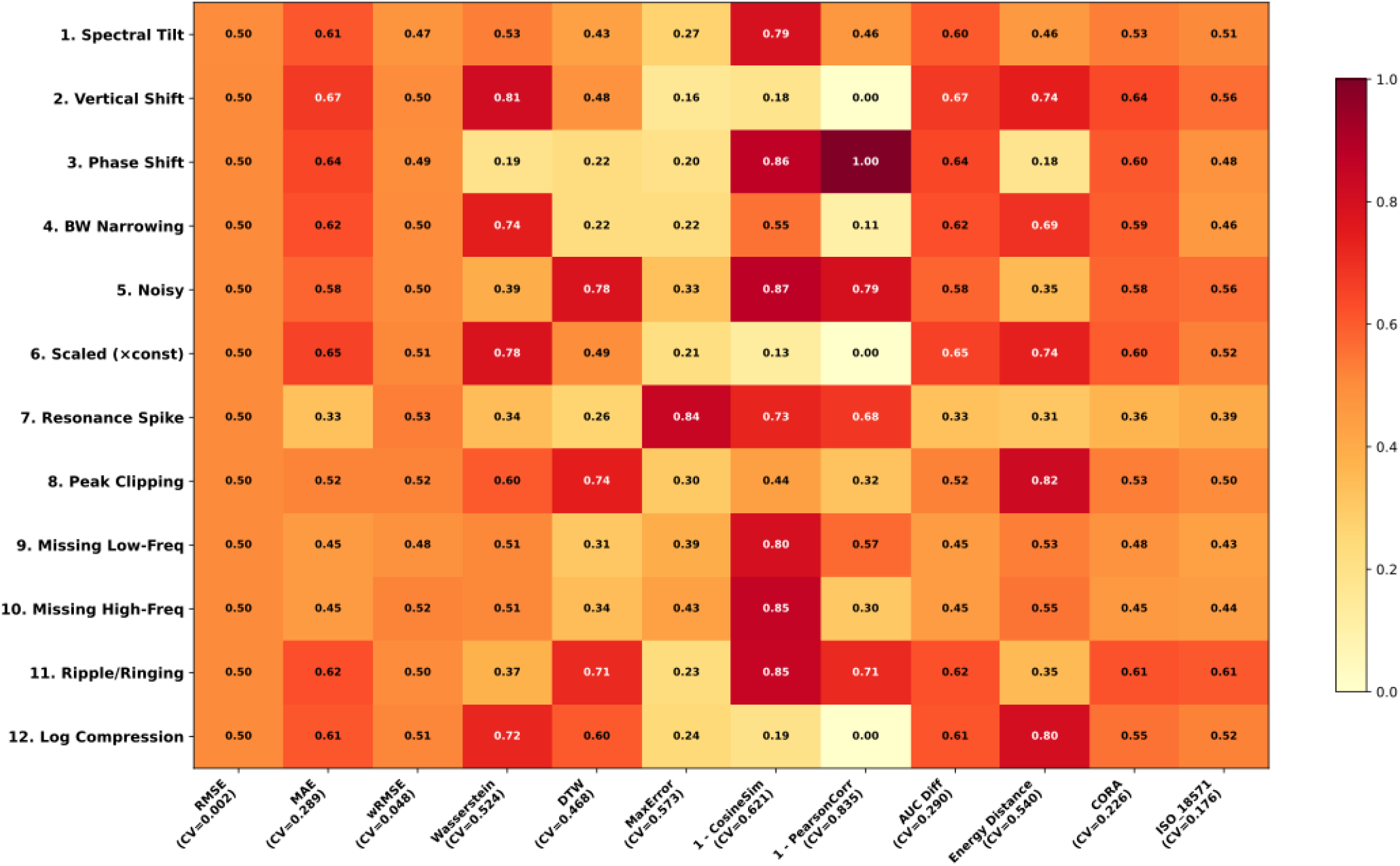
Metric discrimination across controlled spectral distortions with identical RMSE. CV-weighted heatmap of normalized metric responses across twelve controlled error scenarios, all scaled to the same RMSE (0.15). For each metric, scenario penalties were normalized across scenarios and weighted by the metric’s coefficient of variation (CV), which reflects how strongly it differentiates among distortion topologies. Discrimination is reflected by variation within each metric column rather than by the absolute color value itself. Nearly uniform columns indicate limited scenario-specific discrimination, whereas broader within-column spread indicates stronger selectivity across distortion types.

By construction, RMSE remained essentially uniform across scenarios, confirming its inability to separate the iso-RMSE cases. MAE and wRMSE also showed only limited variation, indicating that simple magnitude-based norms provided little additional scenario-specific discrimination. CORA and ISO/TS 18571 showed moderate dispersion across scenarios (CV = 0.39 and 0.65), yet neither provided strong discrimination across the full set of spectral distortion topologies. Several other metrics, however, showed better distortion-specific fingerprints. 1-PearsonCorr spanned the full normalized range from 0.00 to 1.00, with minimal response to vertical shift and maximal response to phase shift, consistent with its insensitivity to uniform level offsets after mean-centering. 1-CosineSim also showed broad selectivity, spanning approximately 0.18 to 0.88, and responded more strongly to global reshaping of the spectrum, such as missing low- or high-frequency content and ripple/ringing, than to simple level shifts. MaxError varied substantially, from approximately 0.20 to 0.83, and emphasized distortions that produced large local pointwise excursions. Among the transport and distribution-based metrics, Wasserstein Distance, DTW, Energy Distance, and AUC Diff also showed some scenario-dependent response patterns, indicating greater sensitivity to feature relocation, nonlinear spectral warping, broad energy redistribution, and cumulative area mismatch than to simple uniform offsets. Overall, the heatmap confirms that no single metric discriminates strongly and consistently across all distortion classes, and that metric behavior depends strongly on the type of spectral mismatch being considered.

### B. Evaluation of Training Dataset Convergence

#### Spectral Evolution and Similarity Trends

Fig. 3 illustrates what is meant by spectral similarity in this analysis. Each colored curve is one simulated FE spectrum, and similarity is evaluated by comparing that individual curve with the mean experimental reference (dashed black line), with the gray band showing experimental variability. In Fig. 3, this comparison is shown only as an example for one representative random seed using MAP derived spectra across training sizes from 100 to 10,000 simulations. The five outputs shown are ear canal (EC) absorbance (*A*_*ec*_), stapes velocity magnitude (|*V*_*st*_|) and phase (∠*V*_*st*_), EC input impedance magnitude (|*Z*_*ec*_|) and phase (∠*Z*_*ec*_). Figs. 4 and 5 extend this same curve to reference comparison across all 10 random seeds and, for each seed, across 11 simulated cases consisting of 1 MAP estimate and 10 posterior samples.

**Fig. 3.**
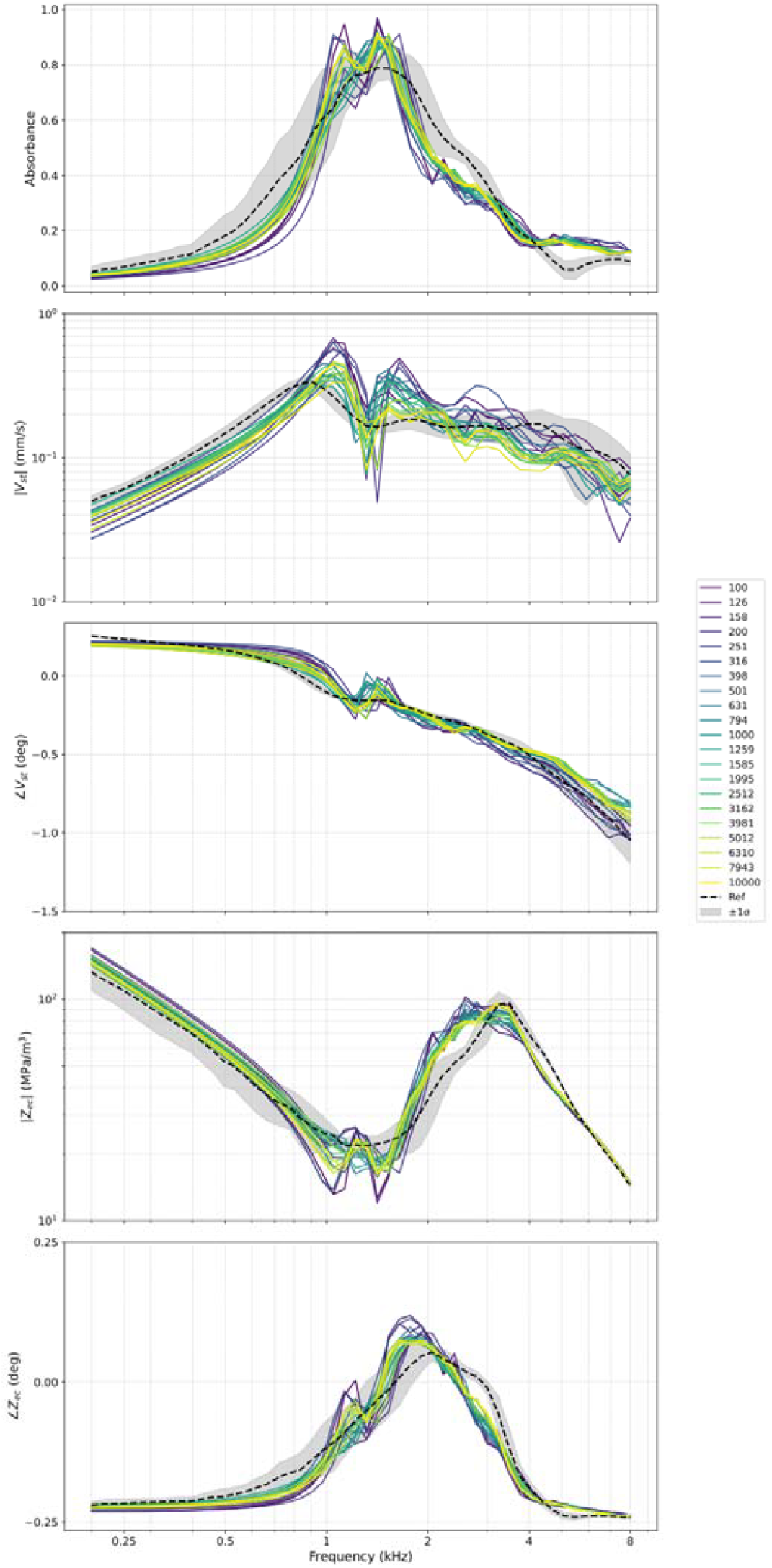
Spectral convergence as a function of training dataset size. Each colored curve is a simulated FE spectrum derived from the MAP estimate at a given training size, N=100 to 10,000, for one representative random seed. Similarity is evaluated by comparing each simulated curve with the mean experimental reference (dashed black line), while the gray band denotes the experimental standard deviation. The five panels show ear canal (EC) absorbance, stapes velocity magnitude and phase, EC input impedance magnitude and phase.

**Fig. 4.**
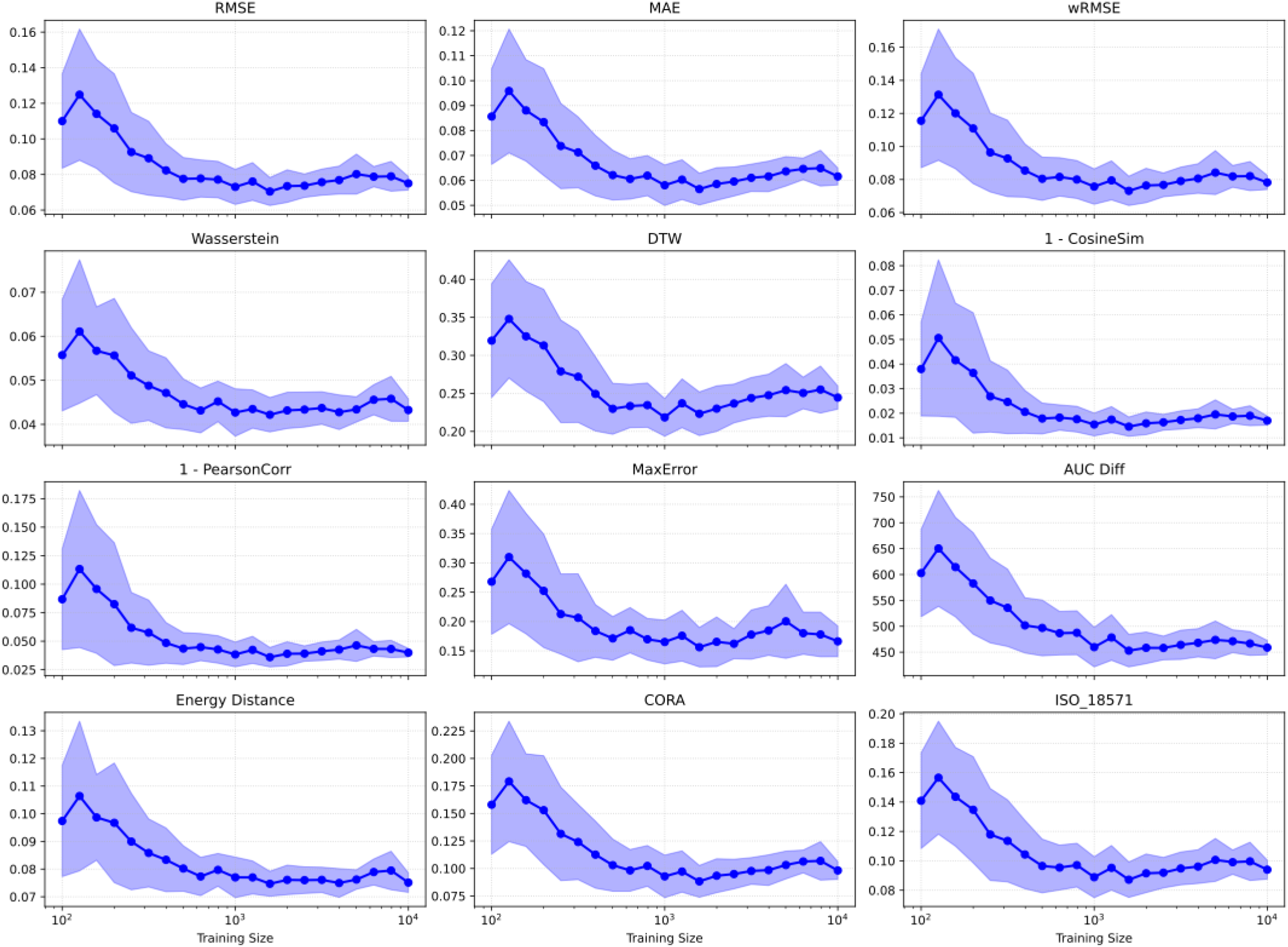
Comparative learning dynamics of individual similarity metrics. Learning curves for twelve similarity metrics computed for the absorbance spectrum, as a function of training dataset size, N=100 to 10,000. For each training size, similarity was summarized across all 10 random seeds and, within each seed, across 1 MAP estimate and 10 posterior samples. Solid lines denote the mean and shaded regions indicate ±1 standard deviation. Most metrics improve rapidly at smaller training sizes and then approach a metric-dependent plateau.

**Fig. 5.**
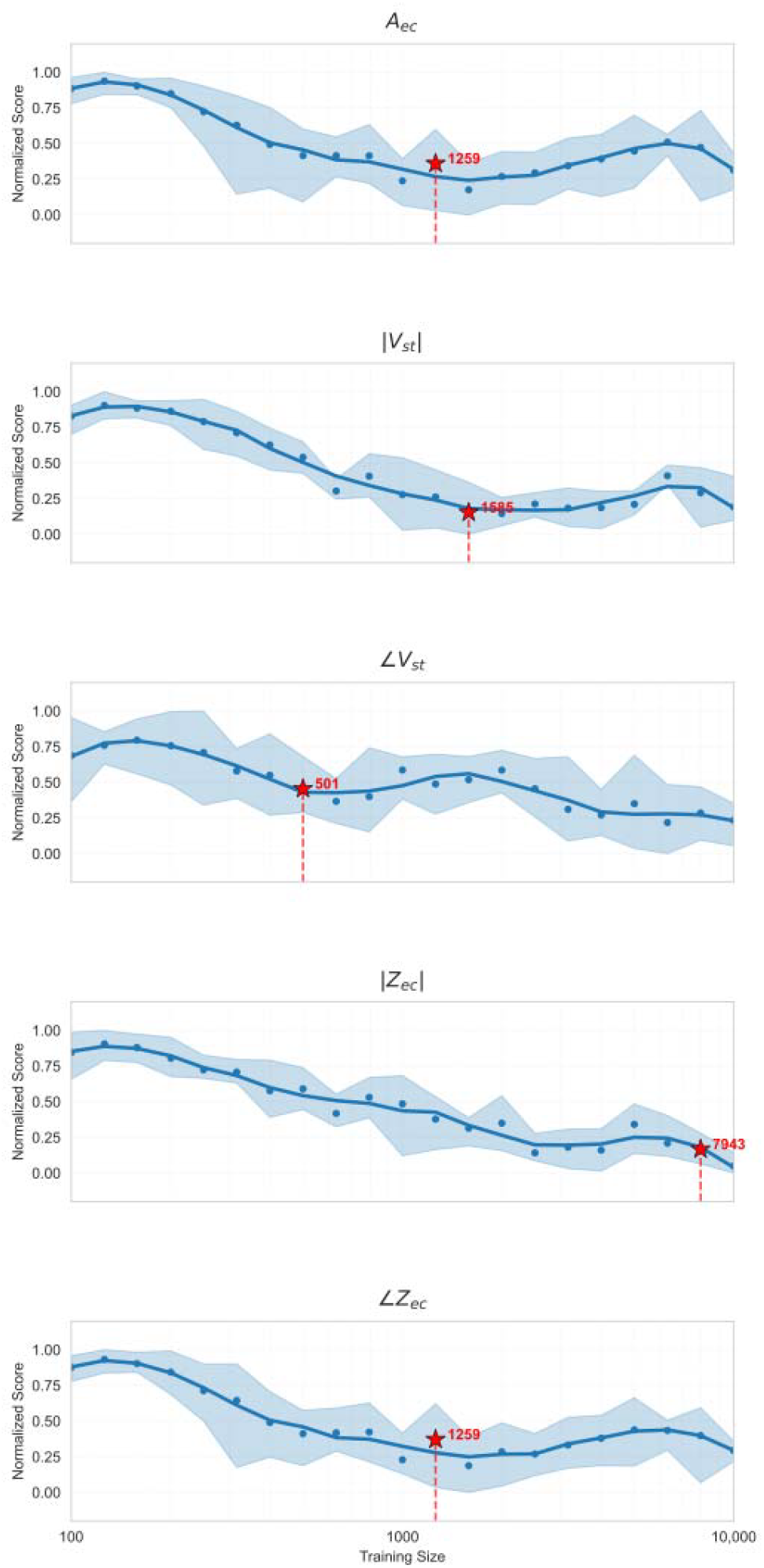
Composite rank aggregation and saturation point identification. Normalized aggregate Borda scores for five spectral measurements as a function of training dataset size. Lower scores indicate better overall agreement. Each point shows the mean across 10 random seeds, where each seed and training size were evaluated using 11 simulated cases consisting of 1 MAP estimate and 10 posterior samples. Solid curves denote the monotonic envelope used for knee-point detection, and shaded bands indicate the 10th to 90th percentile range across seeds. The red star marks the saturation point for each output.

Fig. 4 shows how the individual similarity metrics change with training dataset size for the absorbance spectrum. For each training size, similarity was computed by comparing the experimental reference with the simulated spectra across all 10 random seeds and, within each seed, across 11 simulated cases consisting of 1 MAP estimate and 10 posterior samples. Corresponding results for |*V*_*st*_|, ∠*V*_*st*_, |*Z*_*ec*_|, and ∠*Z*_*ec*_ are provided in the Supplementary Information. Across metrics, most improvement occurred within the first approximately 1,000 simulations, followed by a more gradual plateau. Metric-specific differences were most evident in the convergence tails: shape-based metrics such as 1-CosineSim and 1-PearsonCorr stabilized with relatively low across-seed variability, whereas MaxError showed larger variance and less smooth trajectories, consistent with its sensitivity to localized extremes.

#### Composite Rank Aggregation and Optimal Saturation

Fig. 5 summarizes training-size convergence using a single composite score obtained by rank aggregation of the 12 similarity metrics. Lower normalized Borda scores indicate better overall agreement between the simulated and experimental spectra. Each point represents the mean score at a given training size across 10 random seeds, where each seed was evaluated using 11 simulated cases consisting of 1 MAP estimate and 10 posterior samples. The solid curve shows the monotonic envelope used for knee-point detection, and the shaded band shows the 10th to 90th percentile range across seeds.

The red star marks the saturation point, where improvements shift from rapid to marginal. Using this criterion, saturation occurred at approximately 1,259 simulations for *A*_*ec*_ and ∠*Z*_*ec*_, 501 for ∠*V*_*st*_, and 1,585 for |*V*_*st*_|. In contrast, |*Z*_*ec*_| required at least 7,943 simulations, near the maximum available training size of 10,000, indicating that it was the most data-demanding output for reliable recovery.

### C. Evaluation of Noise Level Robustness

#### Sensitivity of Individual Similarity Metrics to Noise

To evaluate the robustness of SBI performance to stochastic noise inherent in the experimental data, we repeated the SBI training using the full dataset size (10,000 simulations) while adding Gaussian noise to the otherwise deterministic FE outputs at levels scaled to the experimental test-retest standard deviation, σ from 0.1σ to 10σ. Fig. 6 shows the resulting noise-response curves for the 12 similarity metrics computed for the absorbance spectrum, *A*_*ec*_. Corresponding results for |*V*_*st*_|, ∠*V*_*st*_, |*Z*_*ec*_|, and ∠*Z*_*ec*_ are provided in the Supplementary Information. Across metrics, error remained relatively stable at lower noise levels and increased more clearly beyond approximately 1.0σ. Metric-specific differences were evident in both slope and variability: transport and integrated metrics such as Energy Distance and AUC Diff showed comparatively smoother trajectories and narrower across-seed dispersion, whereas metrics sensitive to localized extremes or composite scoring, such as MaxError and CORA, showed larger variance and more irregular responses as noise increased.

**Fig. 6.**
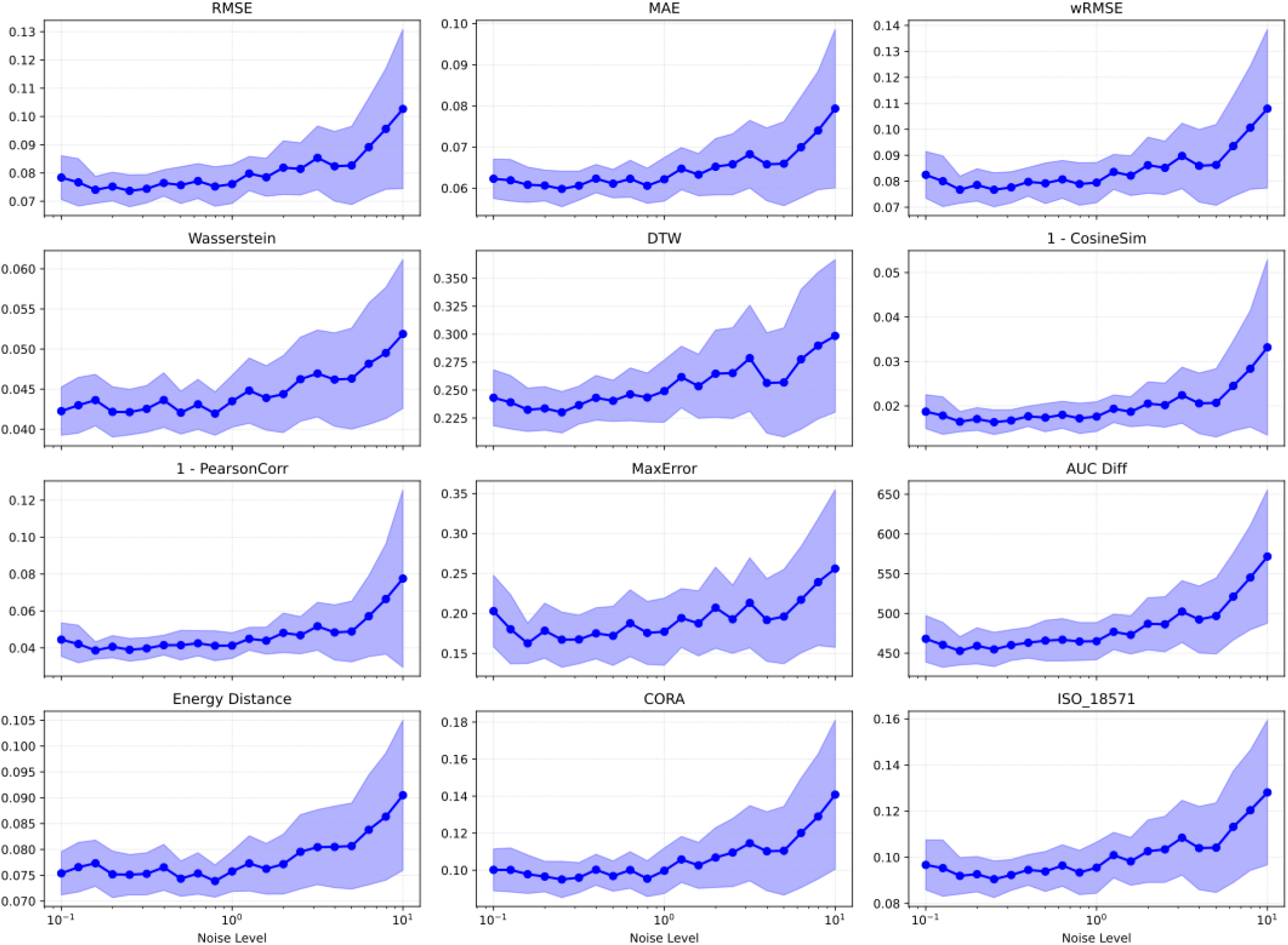
Sensitivity of individual similarity metrics to simulated measurement noise. Noise-response curves for twelve similarity metrics computed for the absorbance spectrum, across simulated noise levels from 0.1σ to 10σ, where σ is the experimental test-retest standard deviation. Solid lines denote the mean across 10 random seeds and shaded regions indicate ±1 standard deviation. Most metrics remain relatively stable at lower noise levels, then show stronger degradation and metric-dependent variability as noise increases.

#### Composite Rank Aggregation and Robustness Limits

Fig. 7 summarizes noise-robustness analysis using a single composite score obtained by rank aggregating the 12 similarity metrics. The normalized Borda score reflects how strongly increasing noise degrades overall agreement between the SBI-derived spectra and the experimental reference, with lower values indicating better agreement. Each point shows the mean score at a given noise level across 10 random seeds, where each seed was evaluated using 11 simulated cases consisting of 1 MAP estimate and 10 posterior samples. The solid curve shows the smoothed trajectory, and the shaded band shows the 10th to 90th percentile range across seeds.

**Fig. 7.**
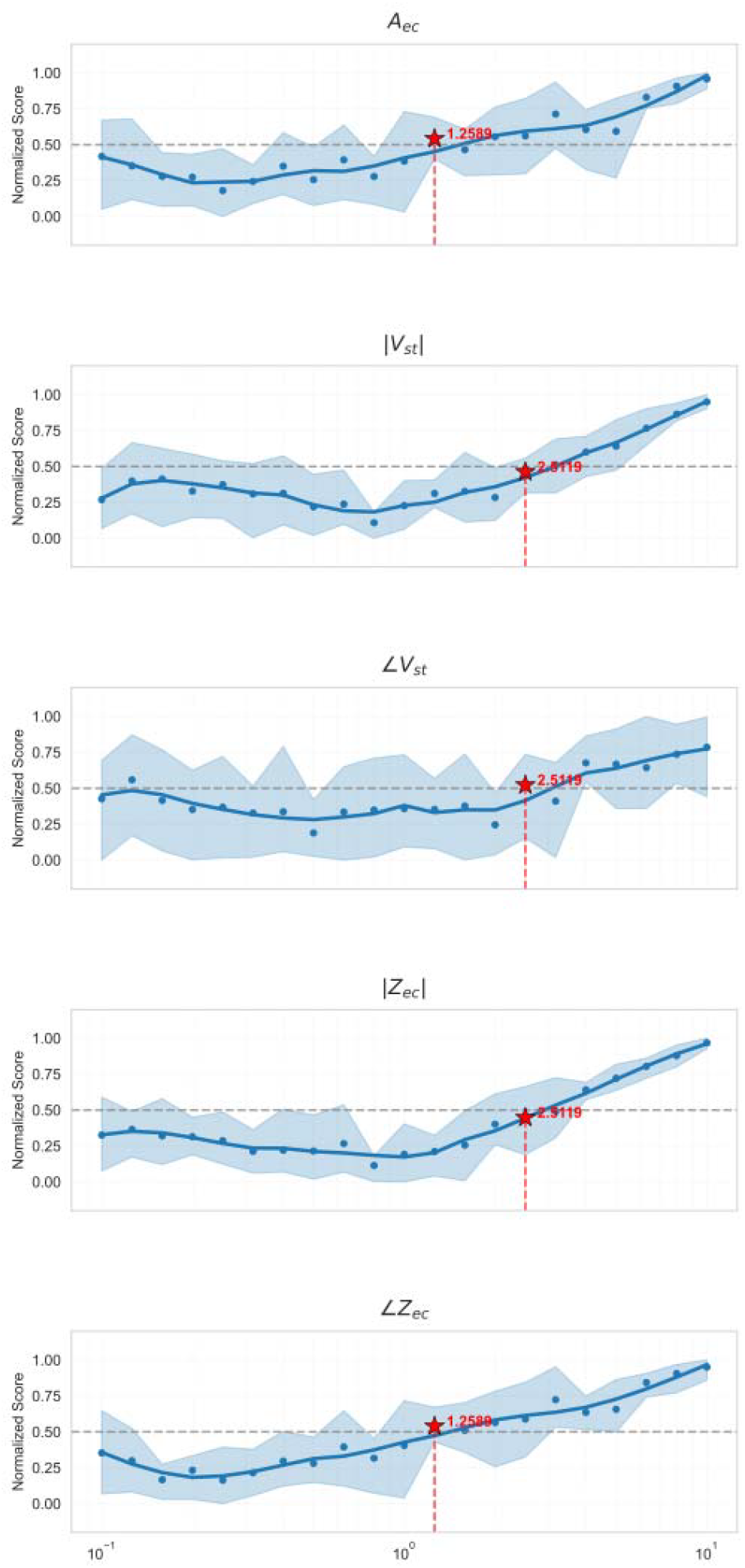
Composite rank aggregation and robustness limit identification. Normalized aggregate Borda scores for five spectral measurements as a function of simulated noise level. Lower scores indicate better overall agreement. Each point shows the mean across 10 random seeds, where each seed and noise level were evaluated using 11 simulated cases consisting of 1 MAP estimate and 10 posterior samples. Solid curves denote the smoothed aggregate trajectory, and shaded bands indicate the 10th to 90th percentile range across seeds. The red star marks the robustness limit, defined as the noise level at which the normalized score reaches 50% of its maximum degradation relative to the noise-free baseline.

The red star marks the robustness limit, defined as the noise level at which the aggregated score reaches 50% of its maximum degradation relative to the noise-free baseline. Noise levels below this point indicate a relatively stable operating range, whereas higher noise levels produce more substantial degradation in SBI performance. Using this criterion, *A*_*ec*_ and ∠*Z*_*ec*_ remained robust up to approximately 1.25σ, whereas |*V*_*st*_|, ∠*V*_*st*_, |*Z*_*ec*_|, and |*Z*_*ec*_| remained robust up to approximately 2.51σ.

## IV. DISCUSSION

### A. Summary of Findings

Spectral biofidelity decisions depend strongly on how similarity is defined and evaluated, particularly when spectra contain multiple resonances and nonuniform distortion patterns (Anderson et al., 2007; Oberkampf et al., 2002). Here, using an SBI-tuned finite-element middle-ear model as a case study (Cranmer et al., 2020; Motallebzadeh et al., 2025; Tejero-Cantero et al., 2020), we evaluated frequency-domain agreement with experimental reference spectra using multiple complementary similarity measures and combined them using a rank-aggregated Borda consensus. This produces a single robust score that is stable across metric choices and stochastic repeats, and is directly usable for comparing conditions in calibration workflows (Dwork et al., 2001; Jurman et al., 2008).

### B. Multi-Metric Consensus to Address Metric Degeneracy

A central outcome of this study is that no single metric can reliably judge spectral biofidelity because distinct distortion topologies can produce similar scalar error values for different underlying deviations (Figs. 1 and 2). This practical metric degeneracy means that resonance shifts, localized spikes, and broadband tilts may be treated as interchangeable by common pointwise or energy-based objectives, despite being mechanistically non-equivalent (Liemohn et al., 2021; Mottershead & Friswell, 1993; Raue et al., 2009).

To reduce these distortion-specific blind spots, we use rank aggregation rather than relying on a single absolute spectral similarity score. Ranking across metrics reduces sensitivity to units, scaling, and dynamic range, and avoids arbitrary cross-metric weighting when combining heterogeneous similarity formulations. The Borda count provides a simple positional consensus that yields stable ordering across conditions and stochastic repeats, converting metric-specific responses into a single decision-ready outcome (Figs. 5 and 7) (Dwork et al., 2001; Jurman et al., 2008; Pihur et al., 2009).

### C. Practical Decision Support and Study Scope

In the context of our SBI implementation, the consensus-based approach is implemented for two practical purposes. First, automated stop criteria, by providing an objective saturation point (elbow or knee point) for neural network training where further training yields diminishing gains in spectral agreement (Figs. 4 and 5). Second, robustness benchmarking, by identifying the noise threshold where the similarity between simulated and experimental spectra is no longer robust across stochastic repeats (Figs. 6 and 7). These decision outputs should be interpreted within the scope of this case study, since convergence and robustness thresholds are expected to vary across specimens, measurement setups, and model structures. Future work should test generalization across additional temporal bones and acquisition conditions, and quantify sensitivity to metric subsets and alternative aggregation rules to characterize how consensus design choices influence conclusions.

### D. Implications for ML Objectives and Physics-Informed Learning

Beyond this case study, high-dimensional spectral comparisons arise across computational biomechanics and ML evaluation whenever agreement must be assessed across multiple outputs and distortion modes. A rank-aggregated multi-metric consensus supports multi-output comparisons without cross-metric scaling or weighting. It also provides a transparent evaluation layer that can be reported with uncertainty across repeats. This aligns with the broader motivation behind composite agreement standards while extending them to frequency-domain spectral biofidelity (Gehre et al., 2009; International Organization for Standardization, 2014; Terven et al., 2025). This also has direct implications for ML loss design: A single norm may be mathematically convenient, but it can miss physically important spectral differences. In contrast, a composite biofidelity measure is more sensitive to distinct physically relevant errors (Goodfellow et al., 2016; Motallebzadeh et al., 2017; Motallebzadeh & Puria, 2022). In physics-informed ML, where the objective must both follow physics and fit/match the data, a composite biofidelity measure can help preserve sensitivity to meaningful spectral distortions across outputs (Bischof & Kraus, 2025), which makes calibration/comparison more interpretable, more defensible, and more physically credible.

## ACKNOWLEDGEMENTS

This work was supported in part by the National Institutes of Health (NIH/NIDCD R21DC020274) and the Canadian Institutes of Health Research (PJT-189955).

## Notes

### Competing Interest Statement

The authors have declared no competing interest.

## REFERENCES

Anderson, A. E., Ellis, B. J., & Weiss, J. A. (2007). Verification, validation and sensitivity studies in computational biomechanics. Computer Methods in Biomechanics and Biomedical Engineering, 10(3), 171–184. 10.1080/10255840601160484

Bischof, R., & Kraus, M. A. (2025). Multi-objective loss balancing for physics-informed deep learning. Computer Methods in Applied Mechanics and Engineering, 439, 117914.

Cranmer, K., Brehmer, J., & Louppe, G. (2020). The frontier of simulation-based inference. Proceedings of the National Academy of Sciences, 117(48), 30055–30062. 10.1073/pnas.1912789117

Dwork, C., Kumar, R., Naor, M., & Sivakumar, D. (2001). Rank aggregation methods for the Web. Proceedings of the 10th International Conference on World Wide Web, 613–622. 10.1145/371920.372165

Gehre, C., Gades, H., & Wernicke, P. (2009). Objective rating of signals using test and simulation responses. 21st International Technical Conference on the Enhanced Safety of Vehicles Conference (ESV), 09–0407. https://www-nrd.nhtsa.dot.gov/pdf/ESV/Proceedings/21/09-0407.pdf

Goodfellow, I., Bengio, Y., & Courville, A. (2016). Deep learning. The MIT press.

International Organization for Standardization. (2014). ISO/TS 18571:2014 Objective rating metric for non-ambiguous signals (Technical Specification ISO/TS 18571:2014). International Organization for Standardization.

Johnson, J., Alahi, A., & Fei-Fei, L. (2016). Perceptual Losses for Real-Time Style Transfer and Super-Resolution. In B. Leibe, J. Matas, N. Sebe, & M. Welling (Eds.), Computer Vision – ECCV 2016 (Vol. 9906, pp. 694–711). Springer International Publishing. 10.1007/978-3-319-46475-6_43

Jurman, G., Merler, S., Barla, A., Paoli, S., Galea, A., & Furlanello, C. (2008). Algebraic stability indicators for ranked lists in molecular profiling. Bioinformatics, 24(2), 258–264.

Liemohn, M. W., Shane, A. D., Azari, A. R., Petersen, A. K., Swiger, B. M., & Mukhopadhyay, A. (2021). RMSE is not enough: Guidelines to robust data-model comparisons for magnetospheric physics. Journal of Atmospheric and Solar-Terrestrial Physics, 218, 105624.

Motallebzadeh, H., Deistler, M., Schönleitner, F. M., Macke, J. H., & Puria, S. (2025). Simulation-based inference for subject-specific tuning of middle ear finite-element models towards personalized objective diagnosis. Scientific Reports, 15(1), 38364. 10.1038/s41598-025-22164-2

Motallebzadeh, H., Maftoon, N., Pitaro, J., Funnell, W. R. J., & Daniel, S. J. (2017). Fluid-structure finite-element modelling and clinical measurement of the wideband acoustic input admittance of the newborn ear canal and middle ear. Journal of the Association for Research in Otolaryngology, 18(5), 671–686.

Motallebzadeh, H., & Puria, S. (2022). Stimulus-frequency otoacoustic emissions and middle-ear pressure gains in a finite-element mouse model. The Journal of the Acoustical Society of America, 152(5), 2769–2780.

Motallebzadeh, H., Soons, J. A., & Puria, S. (2018). Cochlear amplification and tuning depend on the cellular arrangement within the organ of Corti. Proceedings of the National Academy of Sciences, 115(22), 5762–5767.

Mottershead, J. E., & Friswell, M. I. (1993). Model updating in structural dynamics: A survey. Journal of Sound and Vibration, 167(2), 347–375.

Oberkampf, W. L., DeLand, S. M., Rutherford, B. M., Diegert, K. V., & Alvin, K. F. (2002). Error and uncertainty in modeling and simulation. Reliability Engineering & System Safety, 75(3), 333–357.

O’Connor, K. N., Cai, H., & Puria, S. (2017). The effects of varying tympanic-membrane material properties on human middle-ear sound transmission in a three-dimensional finite-element model. The Journal of the Acoustical Society of America, 142(5), 2836–2853.

Pihur, V., Datta, S., & Datta, S. (2009). RankAggreg, an R package for weighted rank aggregation. BMC Bioinformatics, 10(1), 62. 10.1186/1471-2105-10-62

Puria, S. (2003). Measurements of human middle ear forward and reverse acoustics: Implications for otoacoustic emissions. The Journal of the Acoustical Society of America, 113(5), 2773–2789.

Raue, A., Kreutz, C., Maiwald, T., Bachmann, J., Schilling, M., Klingmüller, U., & Timmer, J. (2009). Structural and practical identifiability analysis of partially observed dynamical models by exploiting the profile likelihood. Bioinformatics, 25(15), 1923–1929.

Sackmann, B., Dalhoff, E., & Lauxmann, M. (2019). Model-based hearing diagnostics based on wideband tympanometry measurements utilizing fuzzy arithmetic. Hearing Research, 378, 126–138. 10.1016/j.heares.2019.02.011

Tejero-Cantero, A., Boelts, J., Deistler, M., Lueckmann, J.-M., Durkan, C., Gonçalves, P. J., Greenberg, D. S., & Macke, J. H. (2020). SBI -- A toolkit for simulation-based inference (arXiv:2007.09114). arXiv. http://arxiv.org/abs/2007.09114

Terven, J., Cordova-Esparza, D.-M., Romero-González, J.-A., Ramírez-Pedraza, A., & Chávez-Urbiola, E. A. (2025). A comprehensive survey of loss functions and metrics in deep learning. Artificial Intelligence Review, 58(7), 195. 10.1007/s10462-025-11198-7

Thunert, C. (2017). CORA release 3.6 user’s manual.

Wang, Z., Bovik, A. C., Sheikh, H. R., & Simoncelli, E. P. (2004). Image quality assessment: From error visibility to structural similarity. IEEE Transactions on Image Processing, 13(4), 600–612.

Yang, G., Yang, S., Liu, K., Fang, P., Chen, W., & Xie, L. (2021). Multi-band melgan: Faster waveform generation for high-quality text-to-speech. 2021 IEEE Spoken Language Technology Workshop (SLT), 492–498.

