## Supplementary Information for "Composite Biofidelity: Addressing Metric Degeneracy in Biomechanical Model Validation and Machine Learning Loss Design"

**Table 1.** *Frequency-band weighting scheme used for all similarity metrics.*

| **Frequency Range (Hz)** | **Weight** | **Middle-Ear Mechanical Characteristic** | **Clinical / Functional Relevance** |
| --- | --- | --- | --- |
| **100 – 300** | 0.15 | Stiffness-controlled region; dominated by compliant elements of the tympanic membrane and ossicular joints. | Sensitive to changes in middle-ear compliance such as negative pressure, effusion, or ossicular fixation (O’Connor et al., 2017). |
| **300 – 800** | 0.20 | Transition from stiffness- to inertance-dominated motion as ossicular inertia begins to contribute. | Overlaps lower speech formants; reduced transmission may affect vowel audibility (O’Connor et al., 2017; Ugarteburu et al., 2022) |
| **800 – 2 000** | 0.25 | Primary resonance and peak power-transfer zone of the middle ear. | Highly sensitive to ossicular discontinuity or fixation; corresponds to the main resonance of the ossicular chain (Homma et al., 2009; O’Connor et al., 2017). |
| **2,000 – 5,000** | 0.25 | Mass-controlled transmission region where stiffness effects diminish. | Important for high-frequency consonant perception and detection of mass-related conductive pathologies (Merchant et al., 2019; O’Connor et al., 2017). |
| **5,000 – 8,000** | 0.15 | Extended-high-frequency region; output sensitive to geometric and boundary conditions. | Relevant to extended-frequency immittance measures and early detection of ototoxic or metabolic changes (Caldwell et al., 2006; O’Connor et al., 2017). |

Caldwell, M., Souza, P. E., & Tremblay, K. L. (2006). Effect of Probe Tube Insertion Depth on Spectral Measures of Speech. *Trends in Amplification*, *10*(3), 145–154. https://doi.org/10.1177/1084713806292653

Homma, K., Du, Y., Shimizu, Y., & Puria, S. (2009). Ossicular resonance modes of the human middle ear for bone and air conduction. *The Journal of the Acoustical Society of America*, *125*(2), 968–979.

Merchant, G. R., Siegel, J. H., Neely, S. T., Rosowski, J. J., & Nakajima, H. H. (2019). Effect of Middle-Ear Pathology on High-Frequency Ear Canal Reflectance Measurements in the Frequency and Time Domains. *Journal of the Association for Research in Otolaryngology*, *20*(6), 529–552. https://doi.org/10.1007/s10162-019-00735-1

O’Connor, K. N., Cai, H., & Puria, S. (2017). The effects of varying tympanic-membrane material properties on human middle-ear sound transmission in a three-dimensional finite-element model. *The Journal of the Acoustical Society of America*, *142*(5), 2836–2853.

Ugarteburu, M., Withnell, R. H., Cardoso, L., Carriero, A., & Richter, C.-P. (2022). Mammalian middle ear mechanics: A review. *Frontiers in Bioengineering and Biotechnology*, *10*, 983510.

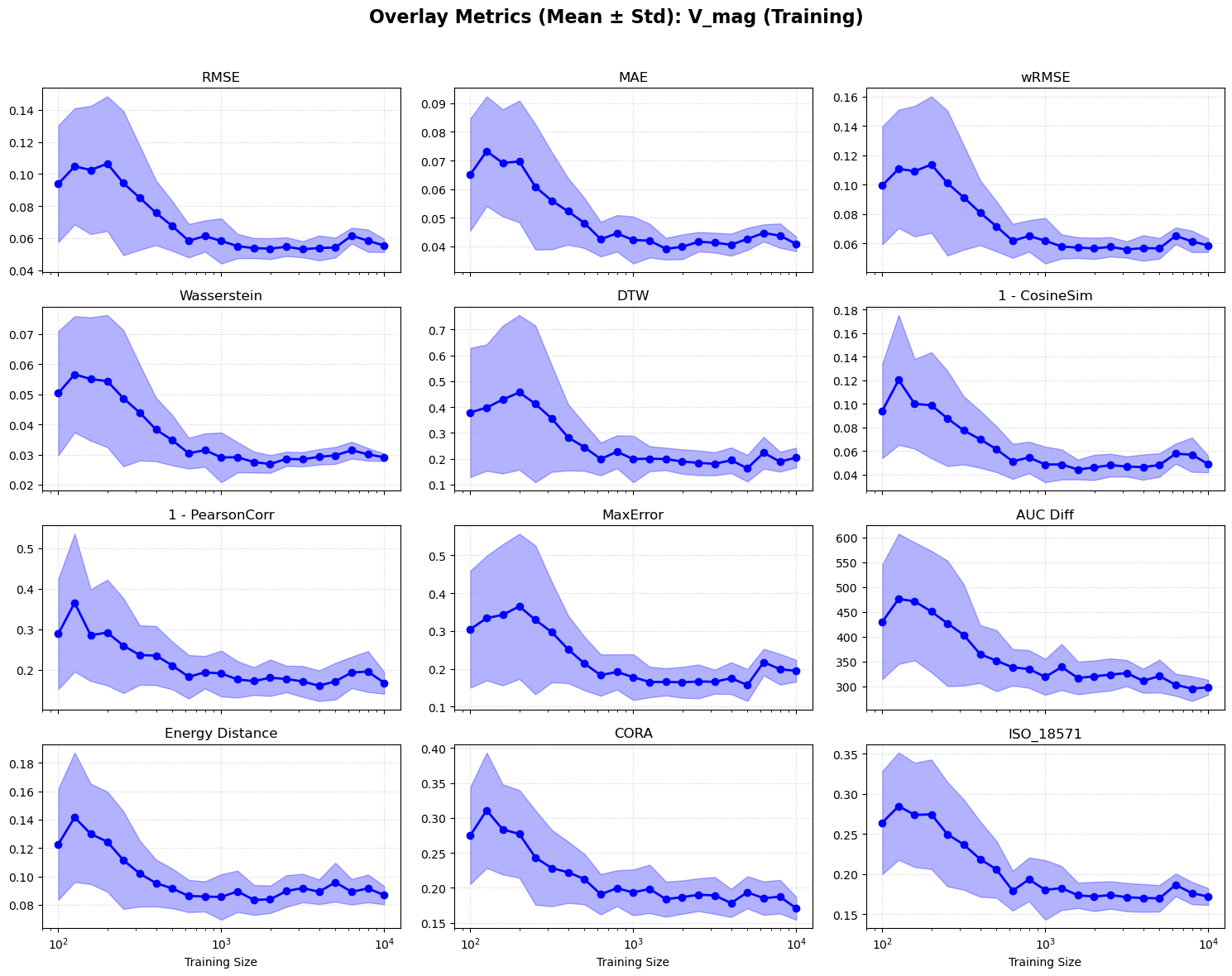

Figure S1. Learning curves for |V_st_|.

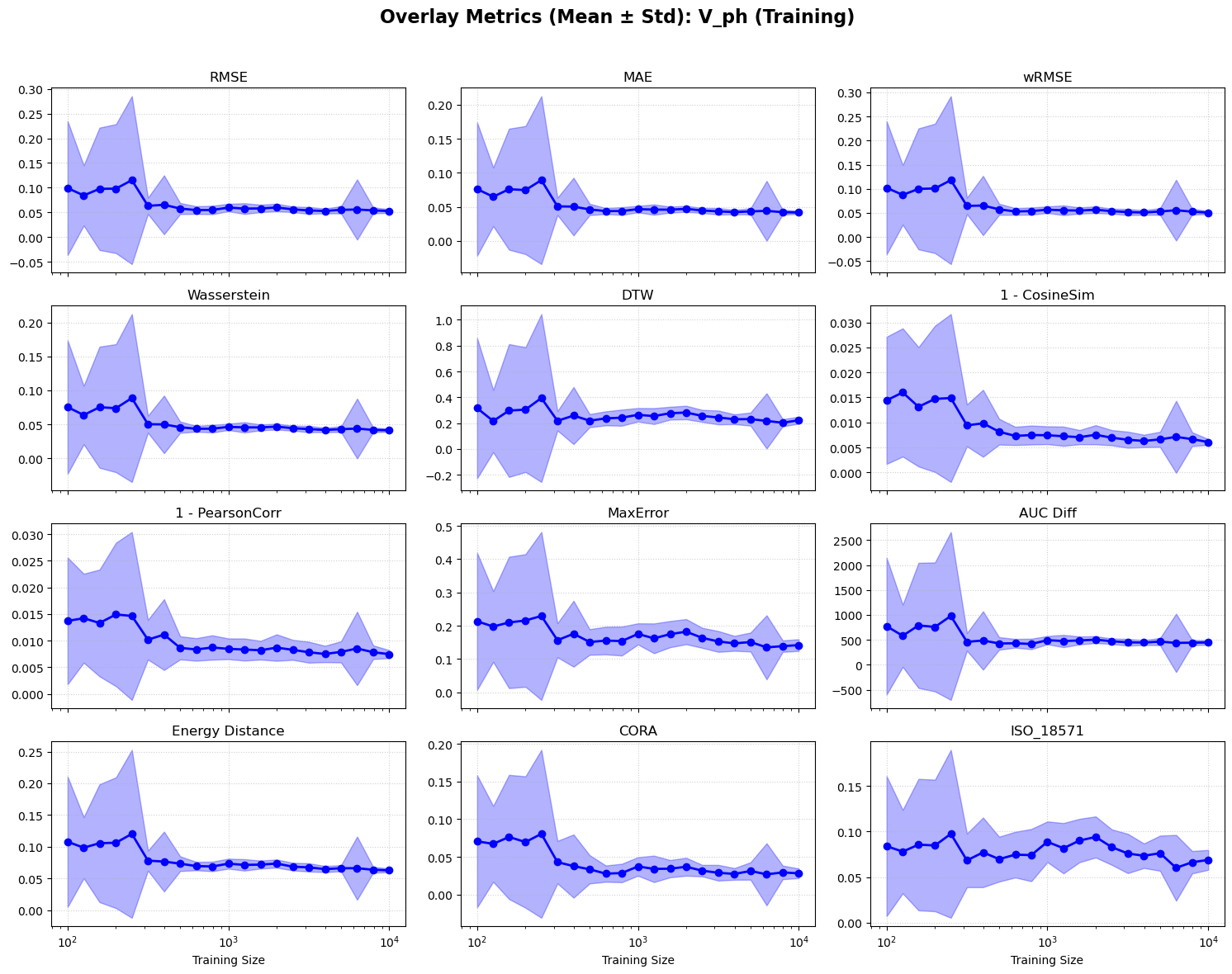

Figure S2. Learning curves for ∠V_st_.

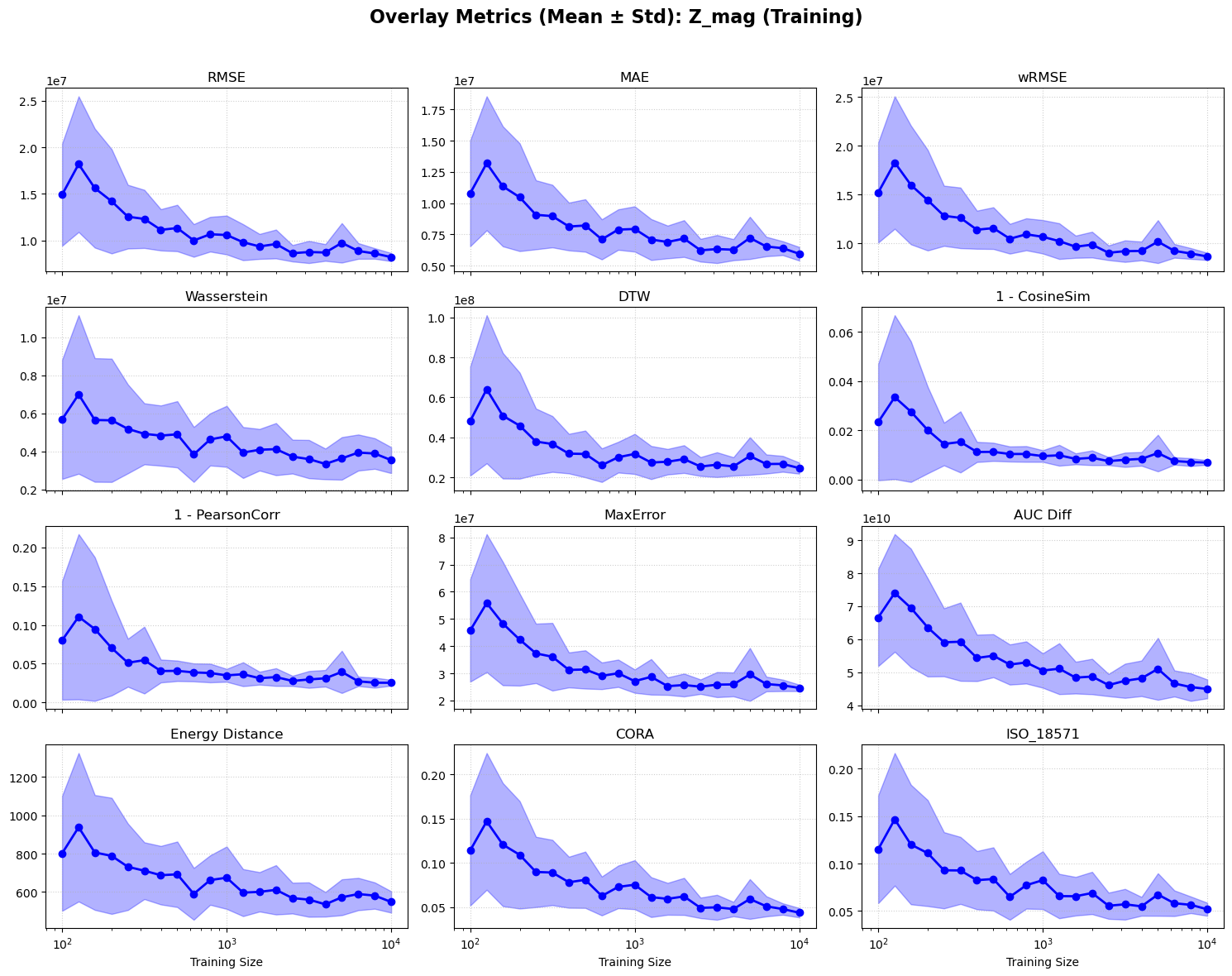

Figure S3. Learning curves for |Z_ec_|.

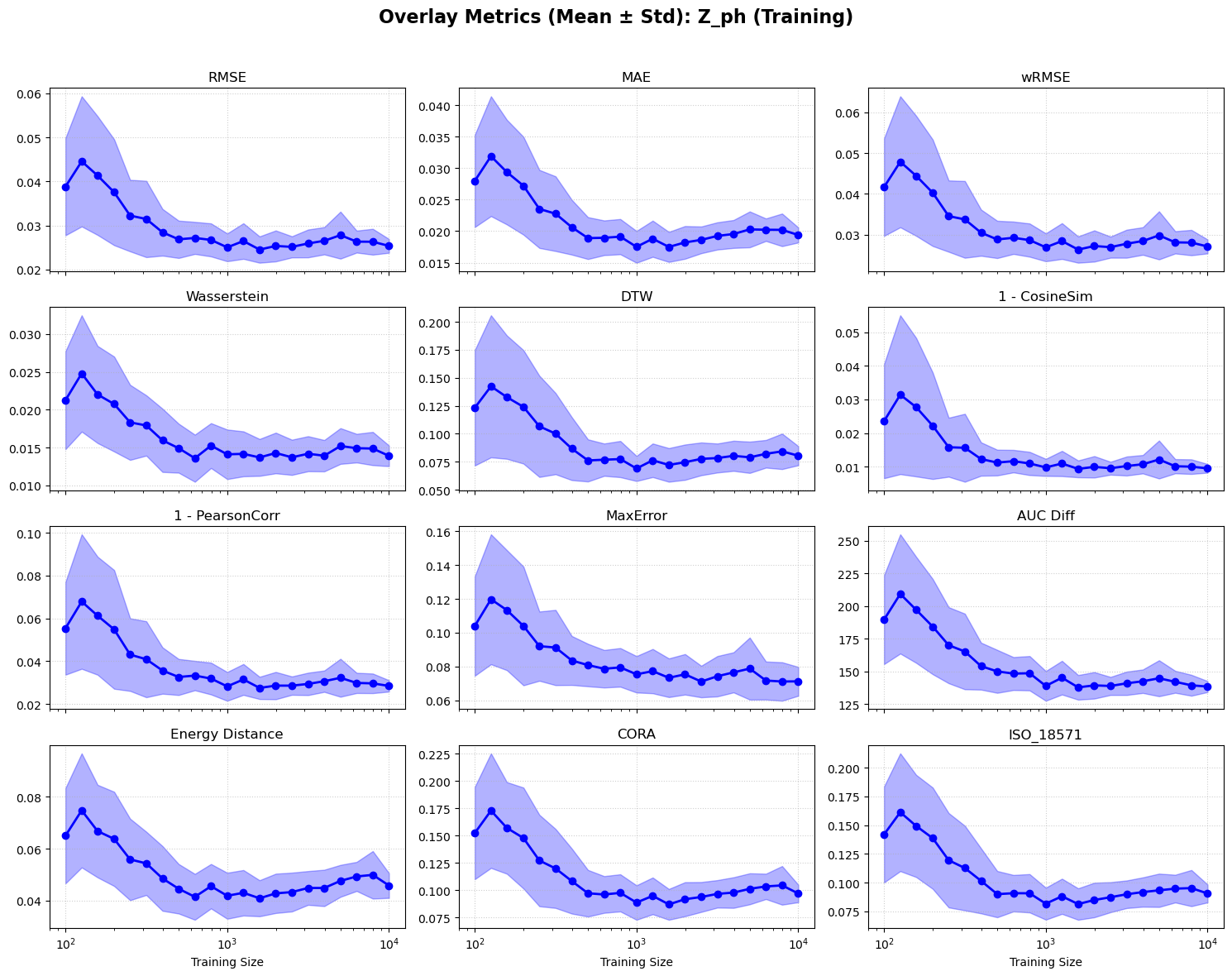

Figure S4. Learning curves for ∠Z_ec_.

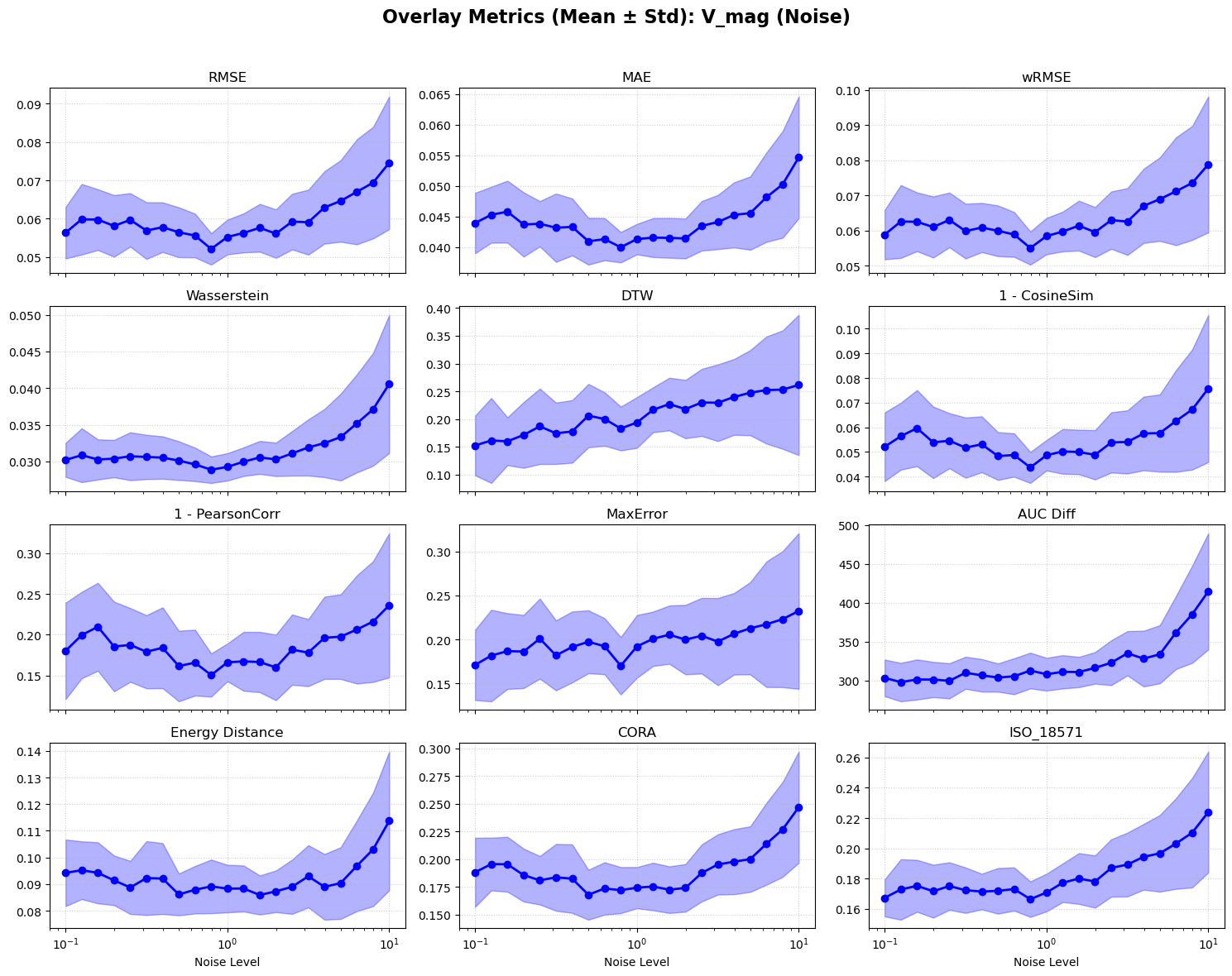

Figure S5. Noise-response curves for |V_st_|.

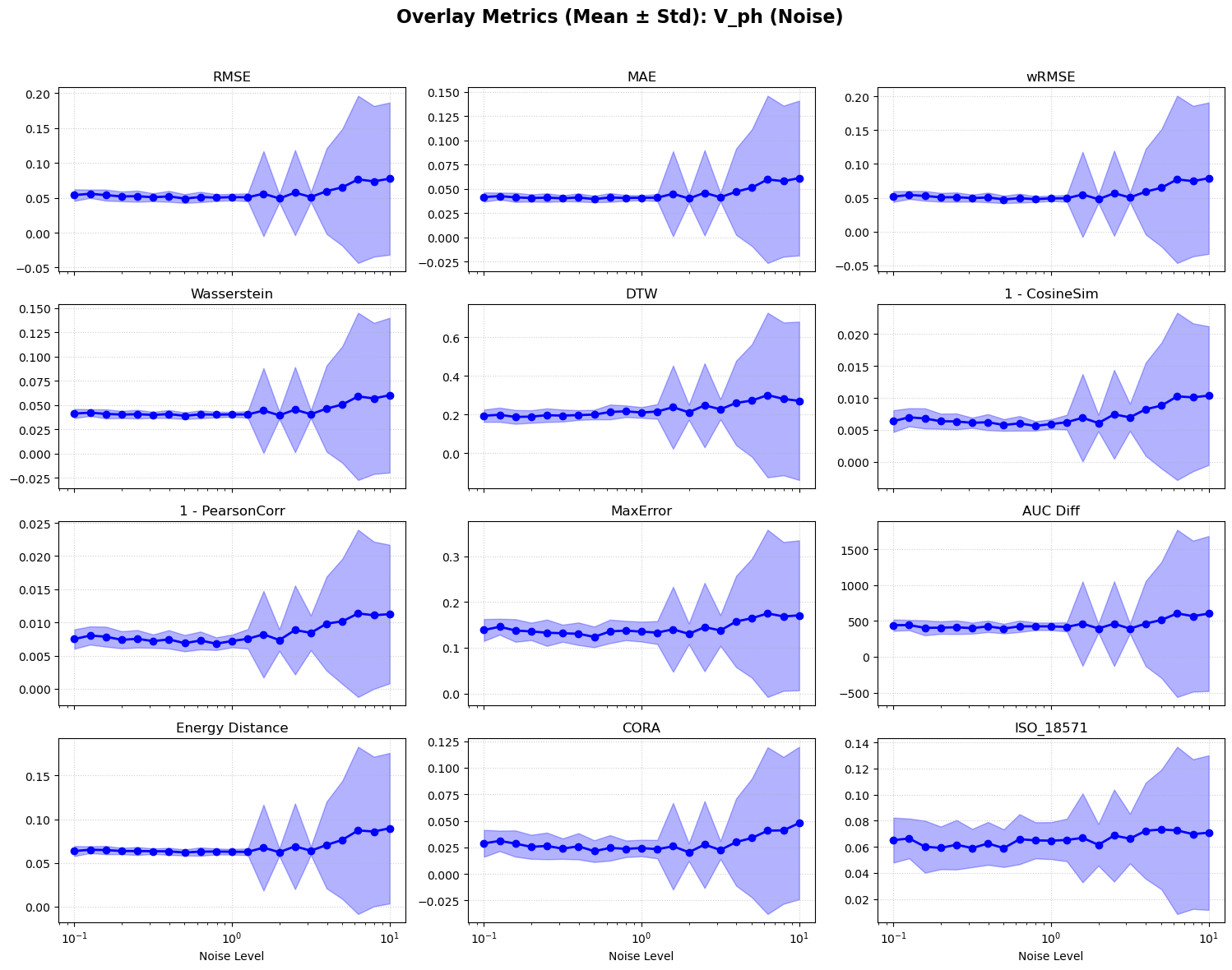

Figure S6. Noise-response curves for ∠V_st_.

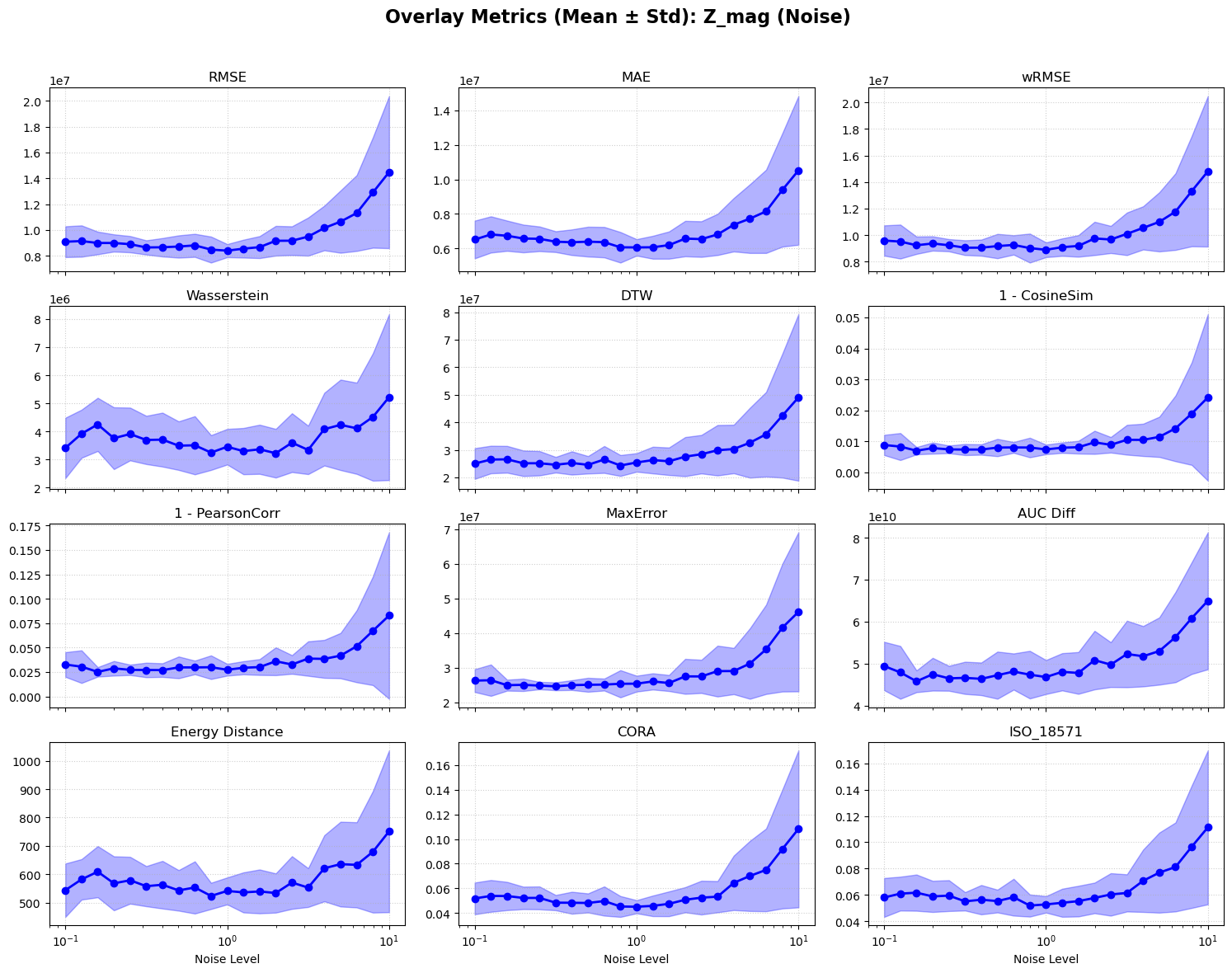

Figure S7. Noise-response curves for |Z_ec_|.

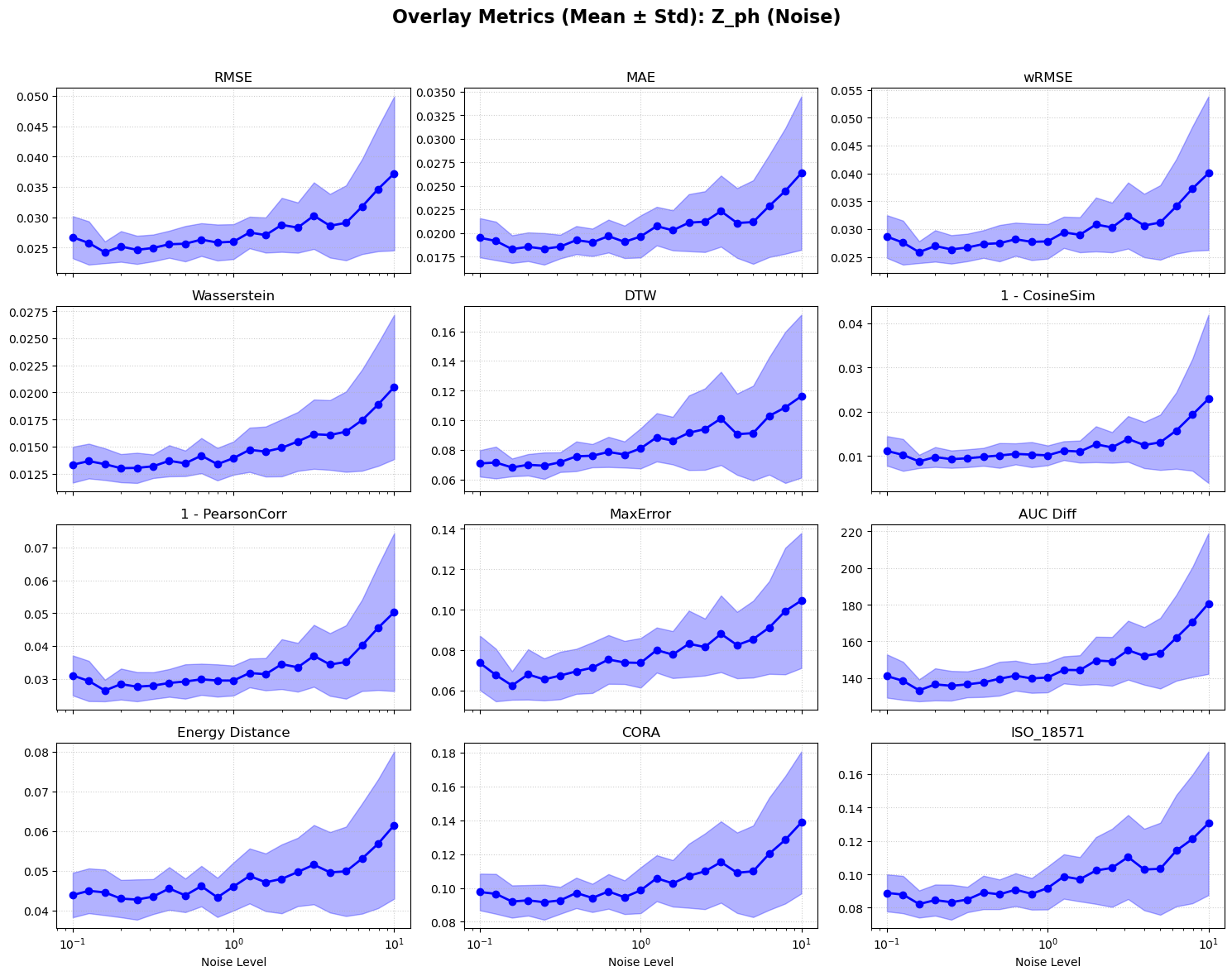

Figure S8. Noise-response curves for ∠Z_ec_.
